# Early microbial intervention reduces diarrhea while improves later milk production in newborn calves

**DOI:** 10.1101/2023.03.30.535013

**Authors:** Yizhao Shen, Yan Li, Tingting Wu, Quanbin Dong, Qiufeng Deng, Lu Liu, Yanfei Guo, Yufeng Cao, Qiufeng Li, Jing Shi, Huayiyang Zou, Yuwen Jiao, Luoyang Ding, Jianguo Li, Yanxia Gao, Shixian Hu, Yifeng Wang, Lianmin Chen

## Abstract

The rumen of neonatal calves is not well-developed and exhibits limited functionality. Therefore, the establishment of intestinal microbiota may play an instrumental role in their health and performance, but it has been rarely explored. Thus, we aim to explore the temporal colonization of the gut microbiome and the potential benefits of early microbial intervention in newborn calves. We followed up on the temporal dynamics of the gut microbiome and plasma metabolome in 36 newborn calves during the first two months of life and established their relationships with their health status and performance. We also evaluated whether microbiota transplantation (MT) could influence their phenotypes by modulating metabolism and its impact on later milk production performance.We showed that the composition and ecological interactions of the gut microbiome are likely to reach maturity one month after birth. Temporal changes in the gut microbiome of newborn calves are widely associated with changes in their physiological statuses, such as growth and fiber digestion. Importantly, we observed that MT reshapes the gut microbiome of newborns by altering the abundance and interaction of *Bacteroides* species, as well as amino acid pathways, such as arginine biosynthesis. Two-year follow-up of those calves further showed that MT improves their later milk production. Notably, MT improves fiber digestion, antioxidant capacity of newborns while reducing diarrhea. MT also contributes to significant changes in the metabolomic landscape, and with putative causal mediation analysis, we suggest that altered gut microbial composition in newborns may influence physiological status through microbial-derived metabolites. The data from the study may help develop strategies to manipulate the gut microbiota during early life, which may be significantly relevant to the health and production of newborn calves.

**HIGHLIGHTS:**

- The gut microbial composition and ecological interaction in newborn calves reach maturity one month after birth
- Temporal shifts in the gut microbiome of newborn calves are associated with phenotypic changes, including growth, digestion, and antioxidative capacity
- Early microbial intervention in newborn calves reduces diarrhea while improving later milk production
- The gut microbial impact on newborn calves is mediated by plasma metabolites

## INTRODUCTION

The colonization and development of the gut microbiota of newborn calves are crucial for the health and performance of the calves later in life (Arshad et al., 2021). This phenomenon is mainly attributed to the fact that these resident microbes support many functions, including the maturation of the immune system (Thaiss et al., 2016), the utilization and modification of nutrients (Nicholson et al., 2012), and the prevention of pathogen colonization (Buffie and Pamer, 2013). Therefore, elucidating the developmental dynamics of gut microbial taxonomy and functionality during early life is important for understanding the relationships between the microbiome and host status and for eventual designing intervention strategies to achieve higher production rates and better health at later stages.

Recent studies have assessed temporal changes in the microbial taxonomic composition of newborn calves during the first week of life (Schwaiger et al., 2020; Song et al., 2018; Takino et al., 2017). For instance, a significant increase in the relative abundance of *Lactobacillus reuteri* was observed during the first week after birth (Schwaiger *et al*., 2020). This fact seems to be very important for calf intestinal health because *L. reuteri* is known to exert bactericidal effects against bacterial pathogens and anti-infective effects against rotaviruses and *Cryptosporidium parvum in vitro* (Schwaiger *et al*., 2020). In addition, comparison of the gut microbial composition between calves (8 weeks after birth) and lactating cows showed that *Bacteroidetes* and *Verrumicrobia* were more abundant in calves, while *Firmicutes*, *Spirochaetes*, *Deinococcusthermus*, *Lentisphaerae*, *Planctomycetes*, and *Chlorofexi* were more abundant in cows (Haley et al., 2020). These observations laid the foundation for targeted mechanistic investigations of the consequences of microbiome colonization for calf health and production.

Nevertheless, several important topics related to the temporal development of the gut microbiome in newborn calves remain unexplored. First, in addition to taxonomy, the functional composition of the gut microbiota can also undergo dynamic changes over time. Microbial functional changes, such as changes in short-chain fatty acid and amino acid metabolic pathways, due to both internal and external disruptions are implicated in the development of immunity and other systems (Visconti et al., 2019; Yang et al., 2020). However, investigations of the temporal dynamics of microbial functionalities of newborn calves are still lacking. Second, the gut microbiome is an ecosystem in which microbes can compete for or exchange nutrients, signaling molecules, or immune evasion mechanisms through complicated ecological interactions that are far from fully understood (Baumler and Sperandio, 2016; Whiteley et al., 2017). These interactions can be identified by co-abundance network analysis and have been shown to be related to human diseases, including obesity and inflammatory bowel diseases (Chen et al., 2020). Investigating temporal changes in microbial interactions during the early life of newborns can enhance our understanding of gut microbial development from an ecological perspective. Third, while the gut microbiome undergoes temporal changes during early life, it is not clear at what time the gut microbiome reaches maturity in calves. This is important because it may indicate a suitable time frame for reshaping the gut microbiome through targeted intervention before maturation. Fourth, a favorable microbiome may promote nutrient utilization and immune responses, but it is not clear whether microbial intervention during early life could improve the digestion, health status of newborn calves, as well as their later milk production performance.

To answer the above questions, we conducted the track dairy cattle study (trackDC) in which 36 newborn calves were randomly assigned into three groups: a control group, a rumen microbiota transplantation group (RMT) and an autoclaved rumen fluid transplantation group (RFT). All the newborn calves from the three groups were followed for 2 months after birth, and intensive phenotype (growth, digestion and fermentation), blood indicator, plasma metabolome and stool metagenomic analyses were conducted (**Figure S1**). In addition, the milk production performance of the calves was recorded during the two-year follow-up period. We not only investigated the temporal development of the gut microbiome at a metagenomic resolution but also evaluated whether MT could influence the phenotypes of newborn calves, including their milk production performance later in life.

## RESULTS

### The Track Dairy Cattle study

To investigate the temporal development of the calf gut microbiome and the potential benefits of early microbial intervention in newborn calves, we collected fecal samples from 36 newborn calves in three groups: a control group, a rumen microbiota transplantation group (RMT), and an autoclaved rumen fluid transplantation group (RFT), as part of the Track Dairy Cattle study (trackDC). Twelve newborn calves were randomly assigned to each group, and we recorded intensive phenotypes, including growth, digestion, ruminal fermentation, and blood measurements at 1, 15, 35, and 56 days after birth (**Figure S1**, **Table S1**). In general, we observed 29 temporal differences and three differences between the groups (FDR < 0.05, **Table S2**). For instance, the digestion rates of acid detergent fiber (ADF) and neutral detergent fiber (NDF) showed temporal differences and were significantly higher in the RMT group than in the other groups, indicating that RMT can promote the digestion of fiber by newborn calves (**Figure 1A-B**, **Table S2**). Moreover, we also observed potential beneficial effects of RMT, as the blood levels of total antioxidant capacity were the highest in the RMT group compared to the other groups (**Figure 1C**, **Table S2**). By comparing the incidence of diarrhea in the three groups, we found that the RMT group had significantly fewer cases of diarrhea (**Figure 1D**). Notably, the two-year follow-up showed that the RMT group had significantly higher milk production than the other groups (**Figure 1E**). Taken together, these results indicate the beneficial effects of early microbial interventions on the health and growth of newborn calves, as well as their later milk production performance.

**Figure 1.**
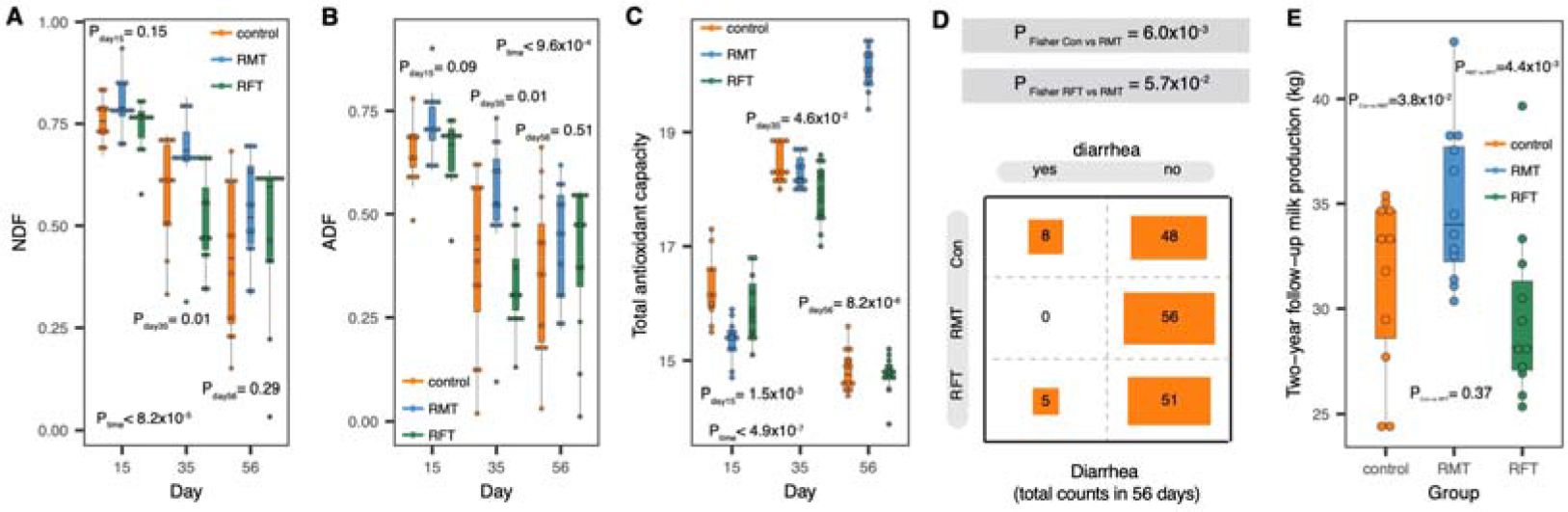
Temporal and microbial transplantation induced variations of phenotypes in newborn calves. A. Temporal changes of neutral detergent fiber (NDF) digestibility. P values from Kruskal test are shown accordingly. **B.** Temporal changes of acid detergent fiber (ADF) digestibility. P values from Kruskal test are shown accordingly. **C.** Temporal changes of plasma total antioxidant capacity. P values from Kruskal test are shown accordingly. **D.** Occurrence of diarrhea between groups in the whole experiential period (56 days). P values from Fisher exact test are shown accordingly. **E.** Differences in the later milk production of those calves during the two-year follow-up. P values from Wilcoxon test are shown accordingly.

### Temporal variations in the gut microbial composition and ecological interaction of newborn calves

To describe the temporal development of the calf gut microbiome, we first evaluated the microbial composition and diversity. A rapid increase in the microbial alpha diversity was observed between 15 and 35 days in all three groups (P_Kruskal test_< 1.1x10^-2^), but there was no significant difference between 35 and 56 days (P_Kruskal test_> 0.05, **Figure S2A**). In addition, dimensionality reduction using the t-distributed stochastic neighbor embedding (t-SNE) algorithm further showed that the microbial composition at Day 15 was significantly different from that at Days 35 and 56 (P_Kruskal test_< 4.0x10^-5^), while no difference was observed between Days 35 and 56 (P_Kruskal test_> 0.05, **Figure S2B**).

For individual microbial species and pathways, we observed that the abundance of 65 out of 125 species (52.0%) and 174 out of 345 pathways (50.4%) were significantly different among the three time points (FDR < 0.05, Kruskal test, **Table S3–4**). Importantly, the temporal development of the gut microbiome was mainly reflected in *Bacteroides* species and the amino acid and nucleotide biosynthesis pathways. In detail, 9 out of 65 differential species were from the genus *Bacteroides*, and 50 out of 174 pathways were related to amino acid and nucleotide biosynthesis (**Table S3–4**). In addition, temporal changes in 80 microbial antibiotic resistance genes and 34 virulence genes were observed (FDR< 0.05, **Table S5**). Interestingly, 57 out of 80 differential microbial antibiotic resistance genes were from *Escherichia coli* (**Table S5**), a widely recognized pathogenic species. Notably, when comparing the mean abundances of microbial species and pathways between time points, we observed that microbial abundances at Day 35 and Day 56 were more similar than those at Day 15 (**Figure S3**). Taken together, these results suggested that the gut microbial diversity and composition of newborn calves likely reached maturity during the first month of life.

In addition to the microbial composition, we further investigated whether microbial interactions, in terms of microbial species and pathway co-abundances, also exhibited differences between different time points in newborn calves. By using the SparCC algorithm (Friedman and Alm, 2012), we established microbial co-abundance relationships in each subgroup separately and identified 2,393 unique species co-abundances and 38,964 pathway co-abundances (P_SparCC_<D0.01, **Table S6–7**). To assess whether microbial co-abundance strengths could be different depending on the time after birth, we assessed to what extent the correlation coefficients were variable across subgroups and observed that on average, 42.3% (ranging from 40.5% to 43.3%) of the species co-abundances and 2.9% (ranging from 0.2% to 7.0%) of the pathway co-abundances showed heterogeneity between different time points (Cochran-Q test, FDR<D0.05, **Figure S4A**, **Table S6–7**). We next summarized the number of differential co-abundances between species from the same genus or from different genera (**Figure S4B**). The genus with the most heterogeneous co-abundances was *Bacteroides*, and many variable co-abundances were observed not only between different *Bacteroides* species but also between *Bacteroides* species and species from other genera such as *Alistipes* (**Figure S4B**). A similar observation was found for the pathway co-abundances, particularly for the nucleotides and amino acid biosynthesis pathways, which showed variability not only within themselves but also with respect to various pathways related to nucleotide biosynthesis (**Figure S4C**). These results indicate that the gut microbiome of newborn calves also undergoes dynamic temporal changes in species interactions after birth, while pathway interactions were relatively stable over time.

### Early microbial transplantation reshapes the gut microbiome composition of newborn calves

As the gut microbial composition likely reaches maturity one month after birth, the accumulating evidence of the importance of the gut microbiota for overall newborn development indicates the need for early modification of the microbiota. Here, we carried out RMT by the oral infusion of fresh ruminal fluid collected from healthy adult cattle. To overcome the potential bias of metabolites in ruminal fluid, which may also influence the development of gut microbiota, we included a group of calves infused with sterilized ruminal fluid. By calculating the inter-calf Bray-Curtis dissimilarity based on the abundances of all the microbial species, we observed that inter-calf dissimilarities within the RMT group were always the lowest among those of all three groups at different time points (P_Kruskal test_ < 1.4x10^-2^, FDR< 0.05, **Figure 2A**). This observation is proof of concept that RMT can reshape the gut microbiome composition of newborn calves, as within group gut taxonomical compositions appear more identical in the RMT group (**Figure 2A**). When comparing individual microbial species and pathways, the relative abundance of 4 species and pathways showed a significant difference at FDR < 0.05 between groups (Kruskal test, **Table S3–4**). Differential microbial species were enriched in the *Bacteroidetes* and *Firmicutes* families, while differential pathways were mostly enriched in nucleotide, amino acid and carbohydrate biosynthesis pathways (FDR < 0.1, **Figure 2B-C**). Notably, differential microbial abundances between groups were mainly observed at Day 15, and most of them were driven by RMT (**Figure 2B-C**), suggesting that RMT had pronounced effects in reshaping the gut microbial composition of newborn calves in the early days of life.

**Figure 2.**
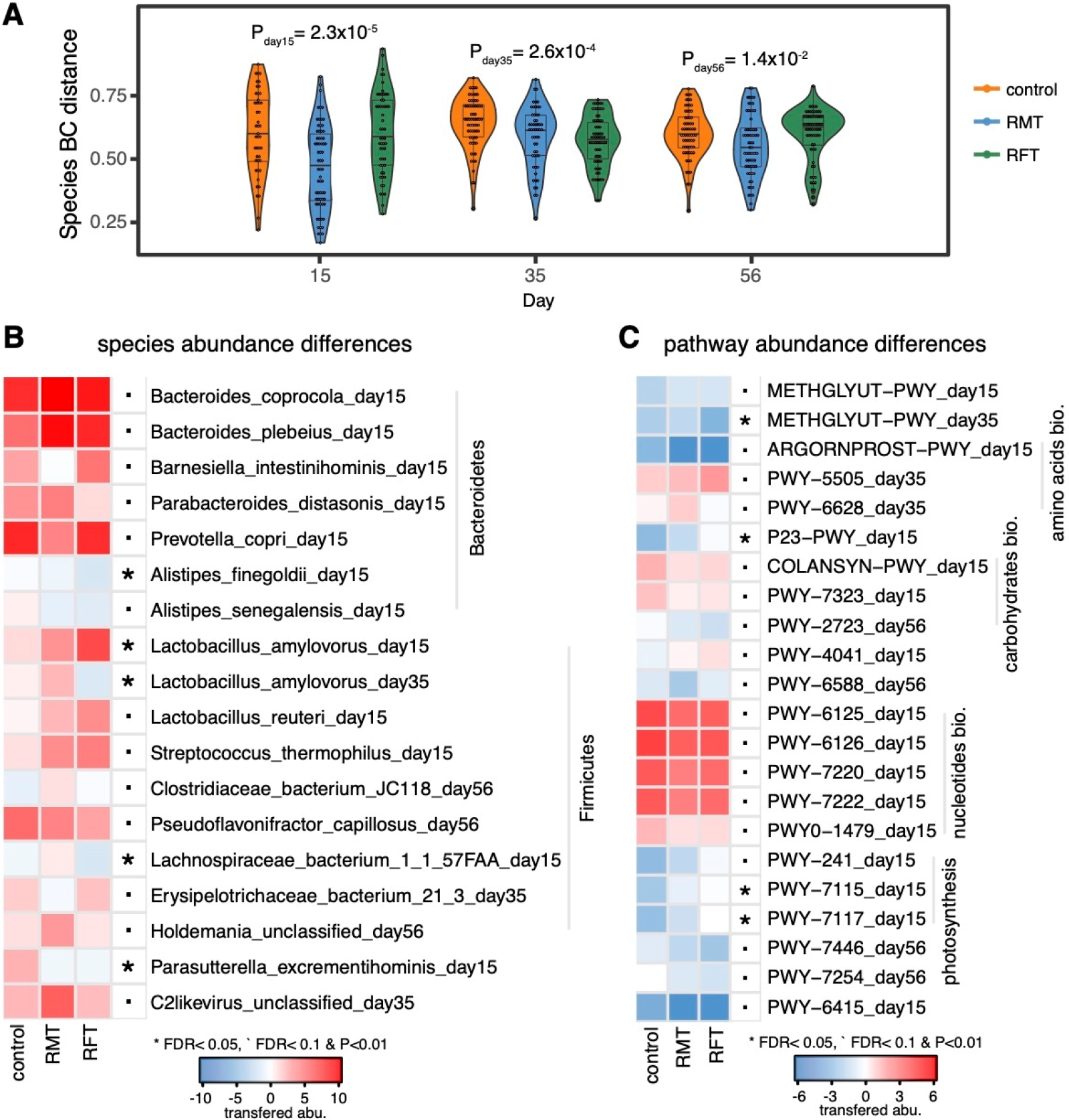
Microbiota transplantation reshapes the gut microbiome composition of newborn calves. A. Within group microbial compositional similarity. The Bray-Curtis (BC) distance represents the dissimilarity of microbial species composition between two samples. P values from Kruskal test are shown. **B.** Differential species abundance between groups. The darkness of color represents the transformed abundance of microbial features. Only results at FDR<0.1 are shown. **C.** Differential pathway abundance between groups. The darkness of color represents the transformed abundance of microbial features.

### Microbial interactions show specificity in the transplantation group

For microbial co-abundances, we have also checked to what extent the correlation coefficients were variable between groups, and the numbers were 40.4% (ranging from 37.1% to 43.5%) and 0.3% (ranging from 0.1% to 0.6%) for species and pathway co-abundances, respectively (Cochran-Q test, FDR<D0.05, **Figure S5A**, **Table S6–7**). These results indicate that microbial interventions can alter the interactions that may potentially contribute to the development of host phenotypes. Interestingly, heterogeneous species co-abundances characterized by comparing different groups were widely distributed in many genera (**Figure S5B**), and these co-bundances were far more complex than those characterized by comparing temporal differences between different time points (**Figure S4B**). However, this was not the case in the pathway co-abundances (**Figure S5C**).

As many of the species and pathway co-abundances showed heterogeneity between groups at different time points, we further analyzed whether those heterogeneous co-abundance relationships were driven by a particular group, i.e., whether the co-abundance strength in one group was very different from those in the other two groups at each time point. In general, we identified 633 and 101 unique group-specific species and pathway co-abundances, respectively (**Table S6–7**). Notably, 248 out of 633 (39.3%) species co-abundances and 42 out of 101 (41.6%) pathway co-abundances showed specificity for the RMT group (**Figure 3A-B**, **Table S6–7**), indicating that RMT can also alter the gut microbiome of newborn calves at the ecological level.

**Figure 3.**
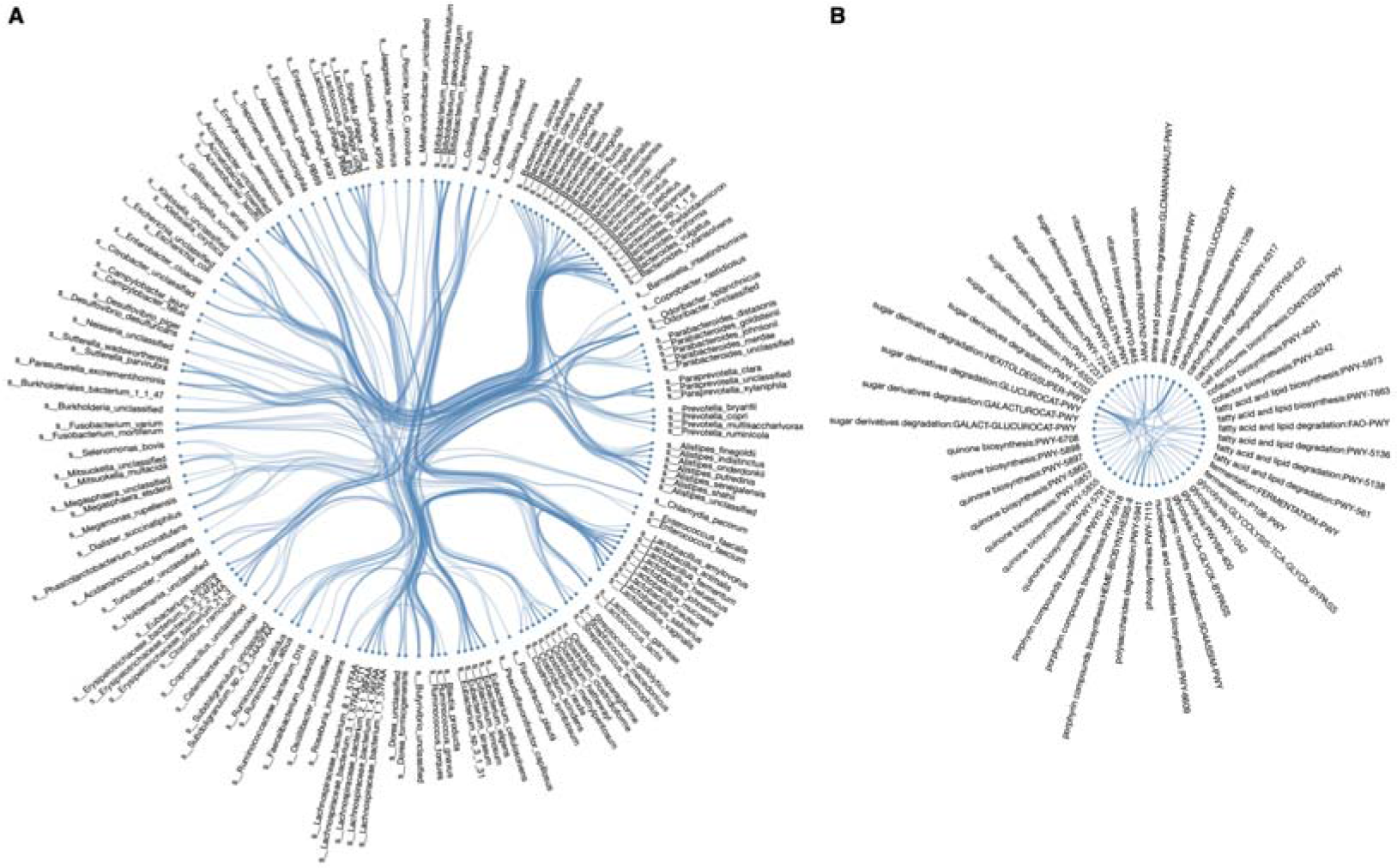
Microbiota transplantation specific species and pathway co-abundances. A. 248 microbial intervention specific species co-abundances. Each dot indicates one species while each line represents a microbial intervention specific correlation between two species. **B.** 42 microbial intervention specific pathway co-abundances. Each dot indicates one pathway while each line represents a microbial intervention specific correlation between two pathways.

Interestingly, we observed a substantial amount of RMT-specific species co-abundances related to *Bacteroides* and *Bifidobacterium* (**Figure 3A**), two common genera that colonized calves early in life (Gacesa et al., 2020; Yassour et al., 2018). For instance, *Bifidobacterium thermophilum* was one of the species with the most RMT-specific co-abundances (10 in total, **Table S6**). *B. thermophilum* constitutes 80% of the infant microbiota and less than 10% of the human adult microbiota, and the presence of *Bifidobacterium* in the gut is often associated with health-promoting effects (Hussein and Singh, 2019). In addition, RMT-specific pathway co-abundances mainly involved sugar derivative degradation and quinone biosynthesis pathways (**Figure 3B**). The sugar derivative degradation pathway-related co-abundances showed specificity for the RMT group, which was reasonable as we observed that the digestion rates of NDF and ADF were relatively higher in the RMT group than in the other two groups (**Figure 1A-B**).

### Microbial changes associated with the phenotypes of newborn calves

To examine the role of gut microbiota colonization in newborn calves, we explored the associations between microbial compositional changes and host phenotypes. To this end, we first calculated the microbial differences between Day 15 and Day 56 and between Day 15 and Day 35. Next, microbial changes were associated with their corresponding phenotypic changes, including changes in growth, digestion, fermentation and blood indicators (**Table S1**). In general, we observed 381 significant associations that involved 79 species, 185 pathways and 16 phenotypes (Spearman correlation, r_absolute_> 0.7, P< 0.01, **Table S8–9**). Most of the microbial associations were related to changes in serum cholesterol, total protein, globulin and albumin levels, suggesting the importance of gut microbial development to newborn calf lipid metabolism and immunity. For instance, changes in the abundance of the microbial saturated fatty acid elongation pathway were associated with changes in serum cholesterol levels (r_Spearman_= -0.85, P= 1.6x10^-3^, **Table S8**). In addition, we observed that changes in serum globulin were associated with 4 prokaryotic ubiquinol biosynthesis pathways (r_Spearman_> 0.79, P< 6.1x10^-3^, **Table S9**). Ubiquinols can promote coenzyme activities to enhance lipophilic antioxidants and thus simulate host immunity (Zhang et al., 2018).

Notably, 77 out of 381 associations were RMT specific (i.e., only showed significance in the RMT group or change in the opposite direction when compared with other groups, r_absolute_> 0.7, P< 0.01, **Table S8–9**), and 54 associations were related to microbial pathways (**Figure 4A-B**). Among those associations, many were related to serum aspartate transaminase (AST), malonaldehyde, total antioxidation capacity, and digestion of fiber (ADF and NDF). This suggests that RMT may potentially influence liver health, energy homeostasis, antioxidation and digestion, as reflected by changes the traits listed above (**Table S6–7**). For example, we observed that an increased abundance of *Megamonas funiformis* may potentially promote the antioxidation capacity of newborn calves (r_Spearman_= 0.77, P= 3.2x10^-3^, **Figure 4C**). In addition, increased microbial arginine biosynthesis was also associated with serum antioxidation capacity (r_Spearman_= 0.80, P= 1.7x10^-3^, **Figure 4D**). Arginine can effectively reduce oxidative stress through the arginine/nitric oxide pathways (Shan et al., 2013). Importantly, we further observed that an increased abundance of *M. funiformis* may contribute to arginine biosynthesis (r_Spearman_= 0.65, P= 2.6x10^-2^, **Figure 4E**), indicating that RMT may increase *M. funiformis* abundance to potentially promote arginine production, which further enhances the antioxidation capacity.

**Figure 4.**
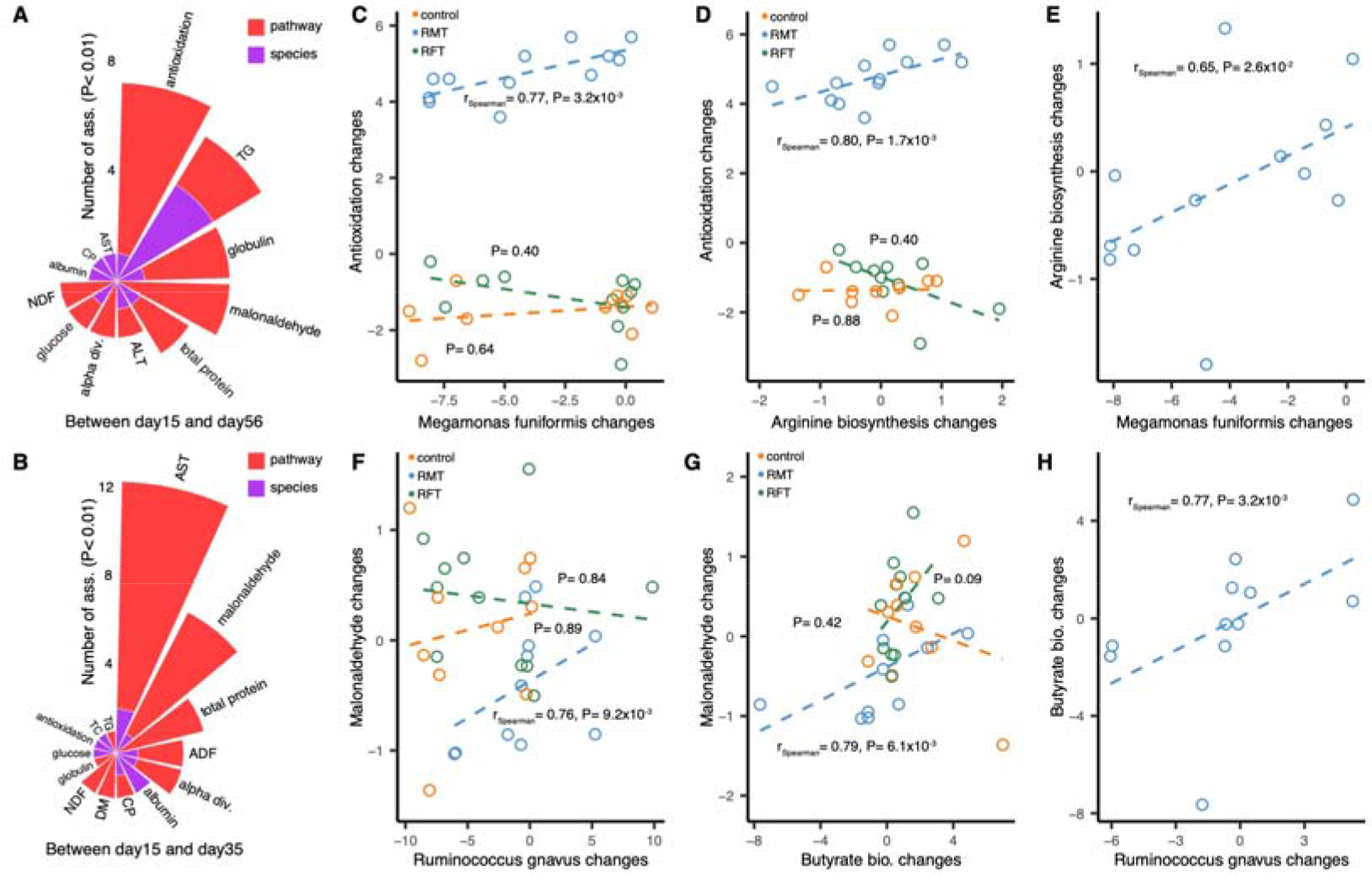
Microbial compositional changes associated with phenotypic changes in newborn calves. **A.** Microbial changes between day 15 and day 56 associated with the corresponding phenotypic changes. **B.** Microbial changes between day 15 and day 35 associated with the corresponding phenotypic changes. **C.** Positive association between antioxidation capacity and *Megamonas funiformis* changes in the microbial intervention group between day 20 and day 56. **D.** Positive association between antioxidation capacity and microbial arginine biosynthesis pathway changes in the microbial intervention between day 15 and day 56. **E.** Positive association between *Megamonas funiformis* and microbial arginine biosynthesis pathway changes in the microbial intervention group between day 15 and day 56. **F.** Positive association between blood malonaldehyde and *Ruminococcus gnavus* changes in the microbial intervention group between day 15 and day 35. **G.** Positive association between blood malonaldehyde and microbial butyrate biosynthesis pathway changes in the microbial intervention group between day 15 and day 35. **H.** Positive association between *Ruminococcus gnavus* and microbial butyrate biosynthesis pathway changes in the microbial intervention group between day 15 and day 35. Spearman correlation is applied to assess the associations between microbial and phenotypic changes.

We also observed that an increased abundance of *Ruminococcus gnavus* was positively associated with changes in the levels of serum malonaldehyde (r_Spearman_= 0.76, P= 9.2x10^-3^, **Figure 4F**), an end product of lipid peroxidation that accumulates in patients with diseases including diabetes (Mahreen et al., 2010). In addition, we observed that increased microbial butyrate biosynthesis was associated with serum malonaldehyde levels (r_Spearman_= 0.79, P= 6.1x10^-3^, **Figure 4G**). Supplementation with butyrate induced a marked shift in superoxide dismutase and catalase activities, along with a decrease in malonaldehyde levels, thereby attenuating oxidative stress (Zhou et al., 2021). Interestingly, we further observed that increased abundance of *R. gnavus* may contribute to butyrate biosynthesis (r_Spearman_= 0.77, P= 3.2x10^-3^, **Figure 4H**), indicating that increased abundance of *R. gnavus* may potentially promote butyrate production in response to the increased amount of malonaldehyde in the RMT group. These results suggest that RMT may potentially promote phenotypes of newborn calves by modulating microbial functionalities.

### Microbial changes associated with plasma metabolites in newborn calves

To further understand the potential mechanisms by which the gut microbiota could influence host physiology, we thought that metabolites are an important class of molecules that are involved in the host-microbe interaction. By profiling plasma levels of 736 metabolites at different time points using untargeted LC-MS (**Table S10**), we observed that the plasma metabolome shifts with time in newborn calves (**Figure S6**) and 50.3% of individual metabolites (370 in total) showed significant differences between groups in at least one time point with FDR <0.05 (Kruskal test, **Table S11**).

We then checked metabolic changes specifically in relation to changes in microbial composition between Day 15 and Day 56 and between Day 15 and Day 35. In total, we observed 17,602 associations between microbial and metabolite changes, and 5,221 of them were RMT specific (Spearman correlation, r_absolute_> 0.7, P< 0.01, **Figure 5A**, **Table S12–13**). Notably, various metabolites that associated with the microbiome are already known to be related to the gut microbiome, including animal essential amino acids, bile acids, organic acids and others (Wang et al., 2022). For instance, increased abundance of microbial L-glutamate biosynthesis pathway (PWY-5505) associated with the increased levels of plasma glutathione (r_RMT_= 0.95, P_RMT_= 4.7x10^-4^, **Figure 5B**), a tripeptide compound consisting of glutamate attached via its side chain to the N-terminus of cysteinyl glycine. Glutathione is an antioxidant that prevent oxidative damage through the reduction of methemoglobin and peroxides (Gaucher et al., 2018). We also observed that increased abundance of microbial L-valine biosynthesis pathway (VALSYN-PWY) associated with the increased levels of plasma N-acetyl-valine (r_RMT_= 0.79, P_RMT_= 9.8x10^-3^, **Figure 5C**). Valine is a branched-chain amino acid that cannot be biosynthesized by animals and plays important roles in insulin resistance and hematopoietic stem cell self-renewal (Cummings et al., 2018; Taya et al., 2016). Taken together, those results suggested that RMT induced temporal changes of the gut microbiome in newborn calves may contribute their metabolic changes.

**Figure 5.**
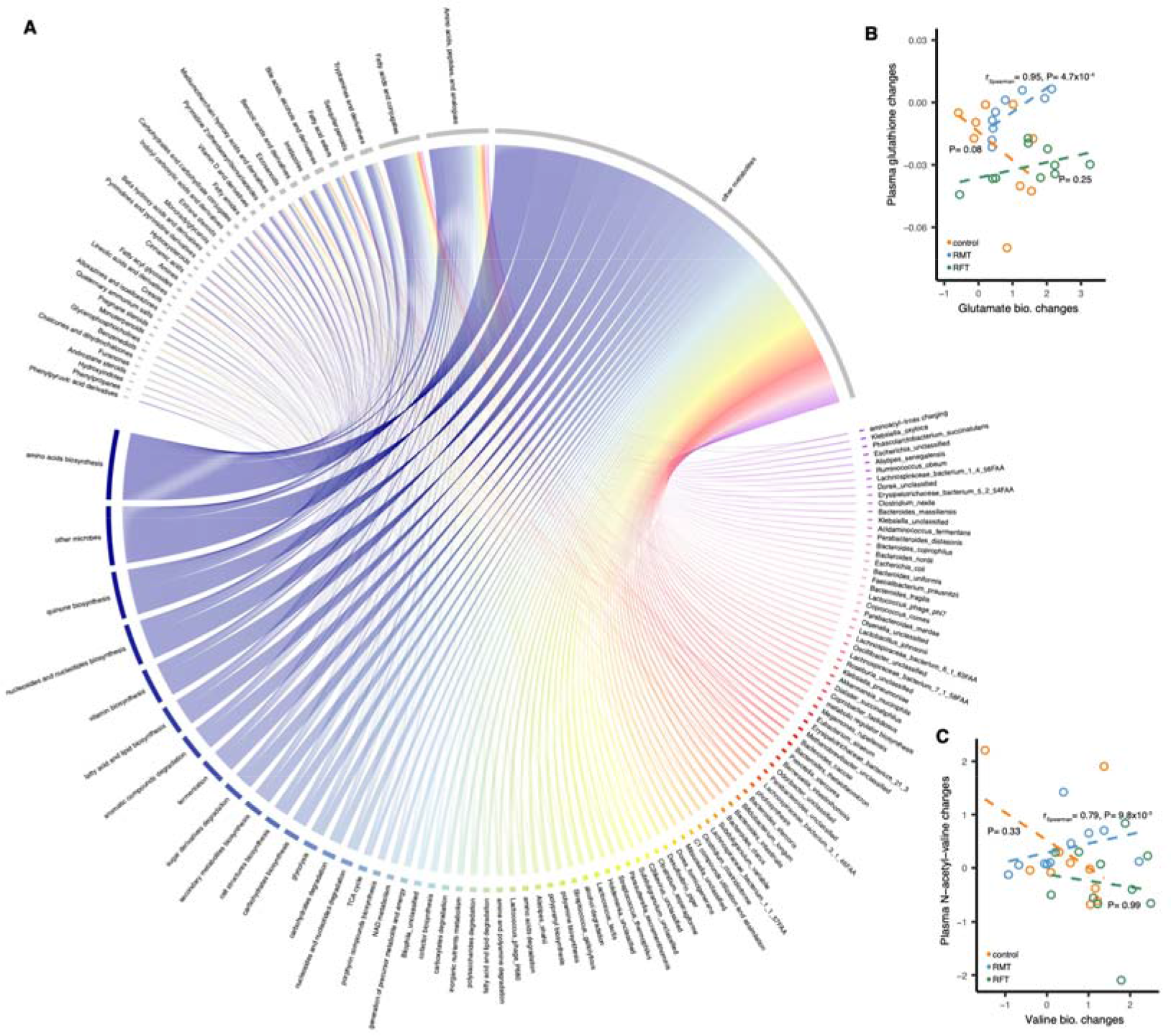
Microbial compositional changes associated with plasma metabolic changes. A. Overview of 5,221 microbial intervention-specific microbe-metabolite associations between day 15 and day 35, day 15 and day 56. The associated microbial factors are colored gray, and the associated metabolites are colored by other colors. **B.** Positive association between plasma glutathione and microbial glutamate biosynthesis pathway changes in the microbial intervention group between day 15 and day 35. **C.** Positive association between plasma N-acetyl-valine and microbial valine biosynthesis pathway changes in the microbial intervention group between day 20 and day 35. Spearman correlation is applied to assess the associations between microbial and phenotypic changes.

### RMT alters phenotypic changes of newborn calves through metabolites

Since microbial changes can be linked to changes in both phenotypes and metabolites in newborn calves, we hypothesized that microbial impacts on host phenotypes may mediate by metabolites. To evaluate whether metabolites can mediate the microbial impact on host phenotypes, we applied mediation analysis focusing on 46 microbial features that are associated with both phenotypic and metabolic changes, which revealed 40 mediation linkages (P_mediation_ < 0.05, **Figure 6A-B**, **Table S14–15**). Those linkages were related to microbial impact on various phenotypes including fiber digestion, antioxidant capacity, lipid and glucose metabolism via a variable category of metabolites (**Figure 6A-B**). For example, we showed that the microbial heme biosynthesis pathway may contribute to an increase in NDF digestion by increasing plasma proline-hydroxyproline levels (P_mediation_ = 0.02, **Figure 6C**). Proline-rich proteins were known as negative regulator that participates in modulating fiber (Xu et al., 2013). For antioxidant capacity, we showed that the microbial purine degradation pathway may contribute to an increase in antioxidant capacity by increasing plasma thymidine levels (P_mediation_ = 0.02, **Figure 6D**). Thymidine catabolism may promote NADPH oxidase-derived reactive oxygen species to induce oxidative stress (Tabata et al., 2018). We also found that a Proteobacteria species *Parasutterella excrementihominis* may contribute to the decrease in plasma triglyceride by increasing plasma androsterone levels (P_mediation_ = 0.02, **Figure 6E**). Androsterone is an effective lipid-lowing agent (Oliver, 1962). In summary, these results suggest that microbial changes induced by RMT may reshape phenotypes of newborn calves through modulating host metabolism.

**Figure 6.**
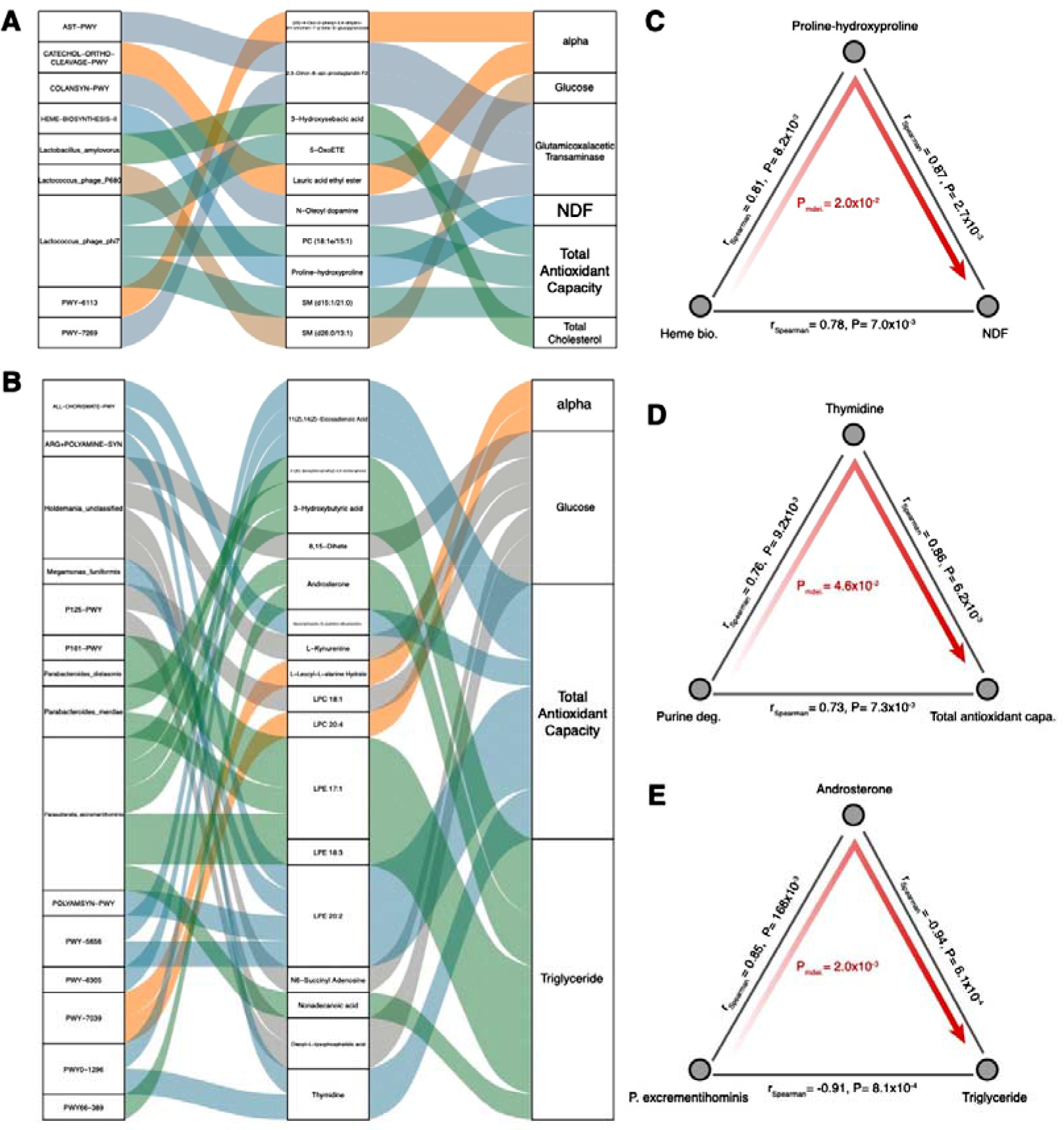
Mediation linkages among the gut microbial, metabolic and phenotypic changes. A. Sankey plot showing the 11 significant mediation effects of plasma metabolites between day 15 and day 35. **B.** Sankey plot showing the 29 significant mediation effects of plasma metabolites between day 15 and day 56. The left panel shows the microbial factors, the middle panel shows the plasma metabolites, and the right panel shows the phenotypes. The curved lines across panels indicate the mediation effects, while the colors correspond to different phenotypes. **C.** Proline-hydroxyproline mediates the effect of microbial heme biosynthesis pathway on NDF digestion. **D.** Thymidine mediates the effect of microbial purine degradation pathway on total antioxidant capacity. **E.** Androsterone mediates the effect of *Parasutterella excrementihominis* on triglyceride.

## DISCUSSION

A diverse microbial population colonizes the mammalian gastrointestinal tract during/after birth, and increasing evidence now suggests that this complex microbiome plays a crucial role in the development of the mucosal immune system and influences newborn health (Arshad *et al*., 2021; Malmuthuge et al., 2015). Recent studies have tracked the temporal changes of the gastrointestinal microbiota in newborn calves during the first several weeks after birth at a 16S rRNA resolution, which is limited on taxonomic composition (Kim et al., 2021; Schwaiger *et al*., 2020; Song *et al*., 2018; Takino *et al*., 2017). However, the key to understanding the importance of microbial development to the host is to investigate whether within-calf microbial differences can be associated with changes in various phenotypes. We therefore systematically characterized the microbial changes at both the taxonomic and functional levels by using fecal metagenomic sequencing data from 36 newborn calves in the trackDC study.

Previous investigations on the temporal development (within 2 months) of the microbial composition of newborn calves with 16S rRNA sequencing have revealed a list of microbial genera that exhibit significant temporal differences, including *Enterococcus*, *Lactobacillus*, *Escherichia*, *Bifidobacterium*, *Clostridium*, etc. (Kim *et al*., 2021; Schwaiger *et al*., 2020; Song *et al*., 2018; Takino *et al*., 2017). Our in-depth metagenomic sequencing of the gut microbiome extends this observation at the species resolution, which enhances the current understanding. In addition, characterization of the temporal changes in microbial pathway abundances further showed that microbial functionalities, such as amino acid, organic acid and carbohydrate metabolism, undergo dynamic changes in newborn calves.

This result was mainly attributed to the fact that microbe colonization of the gastrointestinal tract and their functionalities contribute to changing the ruminant digestive system from a monogastric system to a fully functional foregut rumen fermenter system, with an ability to digest fibrous feed, postweaning. The period from birth to weaning is important for rumen microbial colonization and adaptation. Once the development and maturation of the rumen and the microbiome are complete, it is difficult to permanently manipulate or change the rumen ecosystem due to microbial adaptation and resilience to external mediators (Clemmons et al., 2019). In early life, a favorable microbiome can be implanted via dietary or management interventions and have potentially a long-lasting effect (Arshad *et al*., 2021; Malmuthuge et al., 2019; Palma-Hidalgo et al., 2021). Thus, it is important to determine at what time the gut microbiome of newborn calves reaches maturity. Here, with a longitudinal study design, our data showed that the gut microbiome of newborn calves likely reaches maturity one month after birth, as indicated by the identical microbial diversity, composition and interactions between Day 35 and Day 56 after birth.

Importantly, we further showed that the gut microbiome of newborn calves can be altered by RMT at both taxonomic and functional levels, as indicated by species and pathway abundances, respectively. For instance, we observed a higher abundance of the beneficial species *Parabacteroides distasonis* in the RMT group, and this species can alleviate metabolic dysfunctions by generating succinate and secondary bile acids, which activate the intestinal gluconeogenesis pathway and farnesoid X receptor, in the gut (Wang et al., 2019). In addition, we showed that microbial interactions in terms of co-abundances showed heterogeneity between groups and characterized many RMT-specific species and pathway co-abundances. The diverse microbial communities in the gut make up a complicated ecosystem in which microbes can exchange or compete for nutrients, signaling molecules, or immune evasion mechanisms through ecological interactions that are far from fully understood (Baumler and Sperandio, 2016; Whiteley *et al*., 2017). Our analyses show that microbial alterations by RMT may not be driven solely by differences in abundance level; it may also reflect shifts in microbial interactions that are mirrored in co-abundance analyses. Particularly when applied to metagenomics sequence data, pathway-based co-abundance networks provide further insights into the functional alterations caused by RMT, as many RMT-specific pathway co-abundances have been identified.

Characterization of the temporal changes in the gut microbiome is crucial for understanding the role of the gut microbiome in phenotypic development of newborn calves. By linking microbial changes to phenotypic changes in newborn calves in different groups, we observed thousands of associations between the microbiome, phenotypes and metabolism, including digestion, lipid metabolism, and immunity. Interestingly, many of those associations were only present in the RMT group. For example, RMT may increase *M. funiformis* abundance to potentially promote arginine production, which further enhances the antioxidation capacity. Another example is that increased *R. gnavus* may potentially promote butyrate production in response to the increased amount of malonaldehyde in the RMT group. With mediation analysis by linking microbial, phenotypic and metabolic changes, our analysis suggests that microbial changes induced by RMT may reshape phenotypes of newborn calves through modulating host metabolism. Thus, our longitudinal analysis of microbial association with calf phenotypes and blood indicators revealed functional insights and putative causality of the role of the gut microbiome in newborn calf health status. These observations are of great importance for guiding further studies to develop strategies that may be used to manipulate the early microbiome to improve production and health during the time when newborn calves are most susceptible to enteric disease.

## Limitations of the Study

We acknowledge several limitations in the present study. Firstly, the trackDC study sampled fecal and blood samples from 36 newborn calves during the first two months after birth, making it the first metagenomics and metabolomics-based longitudinal study with the largest sample size to date. However, the sample size is still limited, and replication in independent studies with larger sample sizes may further strengthen our observations and underline their biological significance. Secondly, the reported results are association-based, which means that the underlying causalities and mechanisms of action remain unexplored. Functional studies are thus essential to further reveal the underlying mechanisms of the reported associations. Thirdly, we primarily focused on the gut microbiome of newborn calves. However, given the fact that the temporal development of their rumen microbiome is also important but not easily sampled, further studies linking the gut and rumen microbiomes in newborn calves may provide a more systematic understanding of the importance of the microbiome in newborn calves.

## ACKNOWLEDGEMENTS

We thank the management staff of the trackDC for their supports. This project was funded by the National Natural Science Foundation of China (NSFC) excellent young scientists fund program (overseas) to L.C.; the NSFC surface grant (32270077) to L.C.; the medical expert grant of Jiangsu (2022) to L.C.; the Natural Science Foundation of Jiangsu grant (BK20220709) to L.C.; the Shuang Chuang project of Jiangsu (JSSCBS20221815) to L.C., the NJMU starting grant (303073572NC21 & YNRCZN0301) to L.C.; the China Agriculture Research System of MOF and MARA to Y.G, J.L. and Y.S.; the Hebei Dairy Cattle Innovation Team of Modern Agro-industry Technology Research System (HBCT2018120203) to Y.G. and Y.S.; and the key research and development project of Hebei (20326606D) to Y.G. The funders had no role in the study design, data collection and analysis, decision to publish, or preparation of the manuscript.

## AUTHOR CONTRIBUTIONS

Y.S. and L.C. conceptualized and managed the study. Y.S., Y.L., Q.D, L.L., Y.G., Y.C., Q.L., J.S., H.Z., Y.G., L.D., J.L., Y.G., S.H., Y.W., and L.C. collected the samples and generated the data. Y.S. and L.C. analysed the data. Y.S. and L.C. drafted the manuscript. Y.S., Y.L., T.W., Q.D., Q.D, L.L., Y.G., Y.C., Q.L., J.S., H.Z., Y.G., L.D., J.L., Y.G., S.H., Y.W., and L.C. reviewed and edited the manuscript.

## STAR * METHODS

### KEY RESOURCES TABLE

| REAGENT or RESOURCE | SOURCE | IDENTIFIER |
| --- | --- | --- |
| **Biological Samples** |  |  |
| Fecal samples | This study |  |
| Blood samples | This study |  |
| **Critical Commercial Assays** |  |  |
| QIAamp Fast DNA Stool Kit | QIAGEN | 51604 |
| **Software and Algorithms** |  |  |
| R (3.6.0) | R Foundation | http://www.r-project.org/ |
| MetaPhlan (2.7.2) | (Truong et al., 2015) | PMID: 26418763 |
| HUMAnN2 | (Franzosa et al., 2018) | PMID: 30377376 |
| shortBRED (0.9.5) | (Kaminski et al., 2015) | PMID: 26682918 |
| **Deposited Data** |  |  |
| Metagenomics | ENA | PRJEB42631 |

### RESOURCE AVAILABILITY

### Lead Contact

Further information and requests for resources and reagents should be directed to Lead Contact, Lianmin Chen.

#### Materials Availability

The study did not generate any new reagents or materials.

#### Data and code availability

The metagenomic sequencing data used for the analysis presented in this study are available from the European Nucleotide Archive (ENA) under accession id PRJEB42631. Code used for the analyses is publicly available via: https://github.com/LianminChen/NJMU-LianminChen_Lab/tree/main/trackDC_microbiome

### EXPERIMENTAL MODEL AND SUBJECT DETAILS

#### Animals

The Track Dairy Cattle (trackDC) study is a longitudinal study in northern China that aims to track newborn calves to assess the development of gut microbiota during early life that contributes to cattle health and production. The study was approved by the institutional ethics review board of Hebei Agricultural University (ref. YS19003). In this study, 36 newborn calves were randomly assigned to three groups and followed for two months after birth. The groups included a control group (CON), a rumen microbiota transplantation group (RMT), and a rumen fluid transplantation group (RFT) and intensive data has been collected (**Figure S1**). The newborn calves were trained to feed milk using a bucket and then transferred to individual calf hutches. Starter was provided ad libitum three days after birth and once daily in the morning thereafter. Pasteurized whole milk was fed twice daily at 0800 and 1800 h using a bucket, and the calves were weaned 56 days after birth. RMT and RFT were performed by veterinarians, where the ruminal fluid used in RMT and RFT was collected from a healthy cattle (4-year-old, 600kg, in the dry period,) with a permanent rumen cannula 2 hours after the morning feed. Fresh ruminal fluid was mixed with raw milk and fed to the calves in the RMT group immediately after collection. For the RFT group, the ruminal fluid was autoclaved before feeding. A volume of 50 mL, 80 mL, and 110 mL of ruminal fluid was fed from day 7 to day 11, day 21 to day 25, and day 42 to day 46, respectively. Fecal and blood samples were collected at 15, 35, and 56 days after birth. Besides, the milk production performance has also been recorded during the two-year follow-up.

### METHOD DETAILS

#### Metagenomic data generation and preprocessing

Fecal samples from newborn calves were collected from rectum by stimulation of the anus and stored in liquid nitrogen after well mixed by calf. Aliquots were then made and stored at -80 °C until further processing after transferred to the laboratory. Fecal DNA isolation was performed using the QIAamp Fast DNA Stool Mini Kit (Qiagen, cat.51604). After DNA extraction, fecal DNA was used for library preparation and whole genome shotgun sequencing were performed on the Illumina NovaSeq-6000 platform. From the raw metagenomic sequencing data, low-quality reads were discarded by the sequencing facility, and reads belonging to calf and human contaminations were removed by mapping the data to the reference genomes using Bowtie2 (v.2.1.0) (Langmead et al., 2009; Langmead et al., 2019). After filtering, on average, 36.8 million (sd= 3.6 million) paired reads per sample were obtained for subsequent analysis.

#### Microbial taxonomies

Microbial taxonomic profiles were generated using MetaPhlAn2 (version 2.7.2) (Truong *et al*., 2015). MetaPhlAn2 relies on nearly one million unique clade-specific marker genes identified from approximately 17,000 reference genomes, allowing unambiguous taxonomic assignments, accurate estimation of organismal relative abundance and species-level resolution for bacteria, archaea, eukaryotes and viruses. Microbial species present in more than 10% of the samples were included for further analyses. This yielded a list of 125 species that accounted for 99% of the original species abundance.

#### Microbial pathways

Microbial pathways were determined using HUMAnN2 (Franzosa *et al*., 2018), which maps DNA/RNA reads to a customized database of functionally annotated pangenomes. HUMAnN2 reported the abundances of gene families from the UniProt Reference Clusters (Bateman et al., 2015) (UniRef90), which were further mapped to microbial pathways from the MetaCyc metabolic pathway database (Caspi et al., 2016; Caspi et al., 2018). In total, we identified 345 pathways that were present in at least 10% of samples, retaining 100% of the original functional composition.

#### Microbial antibiotic resistance genes

The abundance of microbial antibiotic resistance genes in metagenomics was determined using shortBRED (version 0.9.5) (Kaminski *et al*., 2015), with markers generated from the CARD database of bacterial antibiotic resistance genes(Jia et al., 2017) (01/11/2018 version). In brief, ShortBRED is a platform for identifying a set of protein sequences from a target database (i.e., ResFinder), clustering them into families, building consensus sequences to represent the families, and then reducing these consensus sequences to a set of unique identifying strings (markers). The platform then searches for these markers in metagenomic data and determines the presence and abundance of the protein families of interest. We classified the abundance of 148 antibiotic resistance genes that were present in at least 10% of the samples.

#### Microbial virulence genes

The abundance of microbial virulence genes was detected using shortBRED (version 0.9.5) (Kaminski *et al*., 2015) and markers generated from virulence factors of the pathogenic bacteria database (VFDB, core dataset of DNA sequences, version: November 2018) (Liu et al., 2019). Then, we classified the abundance of 55 virulence genes that are present in at least 10% of the samples.

#### Growth

The initial body weight was measured immediately after birth, and the final body weight was measured on Day 56 after birth before morning feeding. Body size, including withers height, body length, heart girth, abdominal circumference, and shank circumference, was measured on Day 0 and Day 56 after birth. The starter offered to and refused by each calf were recorded daily during the experimental period. The starter offered was collected by week and refusal was collected daily, pooled by calf weekly. Both offered and refused starter samples were oven-dried at 55 °C for 48 h weekly to determine the dry matter (DM) content. The daily starter DM intake was calculated as the difference between daily starter DM offered and starter DM refused.

#### Digestibility and fecal score

Feed digestibility was determined using acid detergent insoluble ash as an internal marker (Li et al., 2021). Briefly, fecal, starter and milk samples were collected from Day 13 to Day 15, Day 33 to Day 35 d and Day 54 to Day 56, and then, the samples from each calf were pooled, dried at 55 °C for 48 h, and then ground through a 1-mm screen for further analyses. The contents of dry matter (DM, method 930.15) and crude protein (CP, method 996.11) in the starter, milk and fecal samples were determined according to AOAC International. The contents of neutral detergent fiber (NDF) and acid detergent fiber (ADF) in the starter and feces were measured using heat stable α-amylase and sodium sulfite as described by Van Soest et al. (Van Soest et al., 1991). The apparent total tract digestibility was estimated as described by Rice et al. (Rice et al., 2019). The fecal score was monitored and recorded once daily after morning feed on every calf, using a 4-level scoring system, as described by Larson et al. (Larson et al., 1977).

#### Blood biomarkers

Plasma samples were used to analyze the concentrations of blood urea nitrogen (BUN), glucose, total cholesterol and triglycerides, and serum samples were used to analyze the concentrations of total protein, albumin, alkaline phosphatase, aspartate aminotransferase (AST), alanine aminotransferase (ALT), total antioxidant capacity and malonaldehyde. All the blood biomarkers were analyzed using commercial kits from Nanjing Jiancheng Bioengineering Institute (Nanjing, China). The interassay coefficients of variation were lower than 10%, and the intra-assay coefficients of variation were lower than 12%.

#### Ruminal volatile fatty acids and ammonia

Ruminal fluid was collected at Day 56 using an oral stomach tube before morning feeding (Shen et al., 2012). Ruminal pH was measured immediately after collection using a pH mater (Starter 300, Ohaus Instruments Co. Ltd., Shanghai, China). Two subsamples of 5 mL were transferred into 10 mL screw-lid centrifuge tubes after filtering through 4-layer cheesecloth. One subsample was mixed with 1 mL of 25% (wt/vol) HPO_3_ for volatile fatty acid (VFA) analysis, and another subsample was mixed with 1% (wt/vol) H_2_SO_4_ for ammonia analysis. The concentration of ruminal VFAs was measured using gas chromatography (GC-14B, Shimadzu, Japan; 30 m × 0.32 mm × 0.25 mm; column temperature, 110 °C; injector temperature, 180 °C; and detector temperature, 180 °C) (Shen et al., 2019). The concentration of ruminal ammonia was determined as described by Rhine et al. (Rhine et al., 1998).

#### Un-targeted plasma metabolome

Plasma samples resuspended with prechilled 80% methanol. Then the samples were incubated on ice for 5 min and centrifuged at 15,000 g, 4°C for 20 min. The supernatant was injected into the LC-MS/MS system (a ThermoFisher Vanquish UHPLC system coupled with an Orbitrap Q ExactiveTMHF mass spectrometer). The raw data files generated by UHPLC-MS/MS were processed using the Compound Discoverer 3.1 (CD3.1, ThermoFisher) to perform peak alignment, peak picking, and quantitation for each metabolite. The normalized data was used to predict the molecular formula based on additive ions, molecular ion peaks and fragment ions. And then peaks were matched with the mzCloud (https://www.mzcloud.org/) mzVaultand MassList database to obtain the accurate qualitative and relative quantitative results. The annotation of metabolites using the KEGG database (https://www.genome.jp/kegg/pathway.html), HMDB database (https://hmdb.ca/metabolites) and LIPIDMaps database (http://www.lipidmaps.org/).

### QUANTIFICATION AND STATISTICAL ANALYSIS

#### Microbial diversity

The microbial alpha (Shannon index) and beta (Bray-Curtis dissimilarity) diversities were calculated at the species level by using the R (v3.6.0) package *vegan*.

#### Microbial composition dissimilarity

To compare the differences in overall microbial species composition between and within calves in each group at different time points, dimensionality reduction was carried out by using the t-distributed stochastic neighbor embedding (t-SNE) algorithm with the R package *Rtsne*. Microbiome compositional differences between groups and time points were assessed based on one the first and second t-SNE components.

#### Microbial species and pathway co-abundance networks

Microbial species and pathway co-abundance networks were identified by using the SparCC algorithm (Friedman and Alm, 2012). In detail, species composition data from MetaPhlan2 were converted to predicted read counts by multiplying relative abundances by the total sequence counts (Chen *et al*., 2020) and then subjected to SparCC. For pathway analysis, the read counts from HUMAnN2 cells were directly used for SparCC. Significant co-abundance was controlled at the P< 0.01 level using 100 times resampling.

#### Heterogeneity of microbial co-abundances

To assess the variability of networks between groups and different time points, we conducted Cochran-Q tests to assess the heterogeneity of effect sizes and directions for each co-abundance (correlation coefficient generated by SparCC). Here, we treated each subgroup as one study and conducted Cochran’s Q test using the metagen function from the package *meta* (v4.9.5) in R, which calculates the squared difference between individual study effects and the pooled effect using inverse variance weighting (Schwarzer, 2007). For each co-abundance, the P values from the Cochran-Q test were recorded, and co-abundances with significant heterogeneity were controlled at the FDR 0.05 level determined by BH correction.

#### Group specific microbial co-abundances

For heterogeneous co-abundances (Cochran-Q test FDR< 0.05), we further assessed whether these relationships showed group specificity, i.e., whether the effect size of co-abundance (SparCC correlation coefficient) in one group was very different from that in the other two. We adopted interquartile ranges based on the outlier detection method (Barbato et al., 2011). The interquartile range (IQR) was calculated based on the effect size of co-abundances in each group, and we assessed whether the smallest or largest effect size fell outside of Q1-0.75IQR or Q3+0.75IQR. If only one met the condition, we called this co-abundance specific and assigned it to the corresponding group.

#### Differential phenotypic, microbial and metabolic features

The relative abundances of both species and pathway datasets were centered log-ratio transformed, followed by inverse-rank transformation, before subsequent analysis (Aitchison, 1982). No transformation was applied to phenotypic data. The ranked-based Kruskal test was then applied to assess 1) whether features within certain groups showed differences between different time points and 2) whether features within certain time points showed differences between different groups. The false discovery rate (FDR) was calculated by using the Benjamini-Hochberg (BH) method (Benjamini and Hochberg, 1995).

#### Microbial changes linked to phenotypic and metabolic changes

Since we observed that microbial differences between Day 35 and Day 56 were minor, we only calculated microbial and phenotypic changes between Day 15 and Day 35 and between Days 15 and 56. We further linked microbial changes to host phenotypic and metabolic changes with Spearman correlation. Association with P< 0.01 were considered significant.

#### Mediation linkage inference

For phenotypic and metabolic associations to the same microbial feature, we first checked whether the human phenotype was associated with the metabolite using Spearman correlation (P<0.01). Next, mediation analysis was carried out using the mediate function from the R package mediation (version 4.5.0) to infer the causal role of the microbiome in contributing to the human phenotype through metabolites.

---

**Figure S1.**
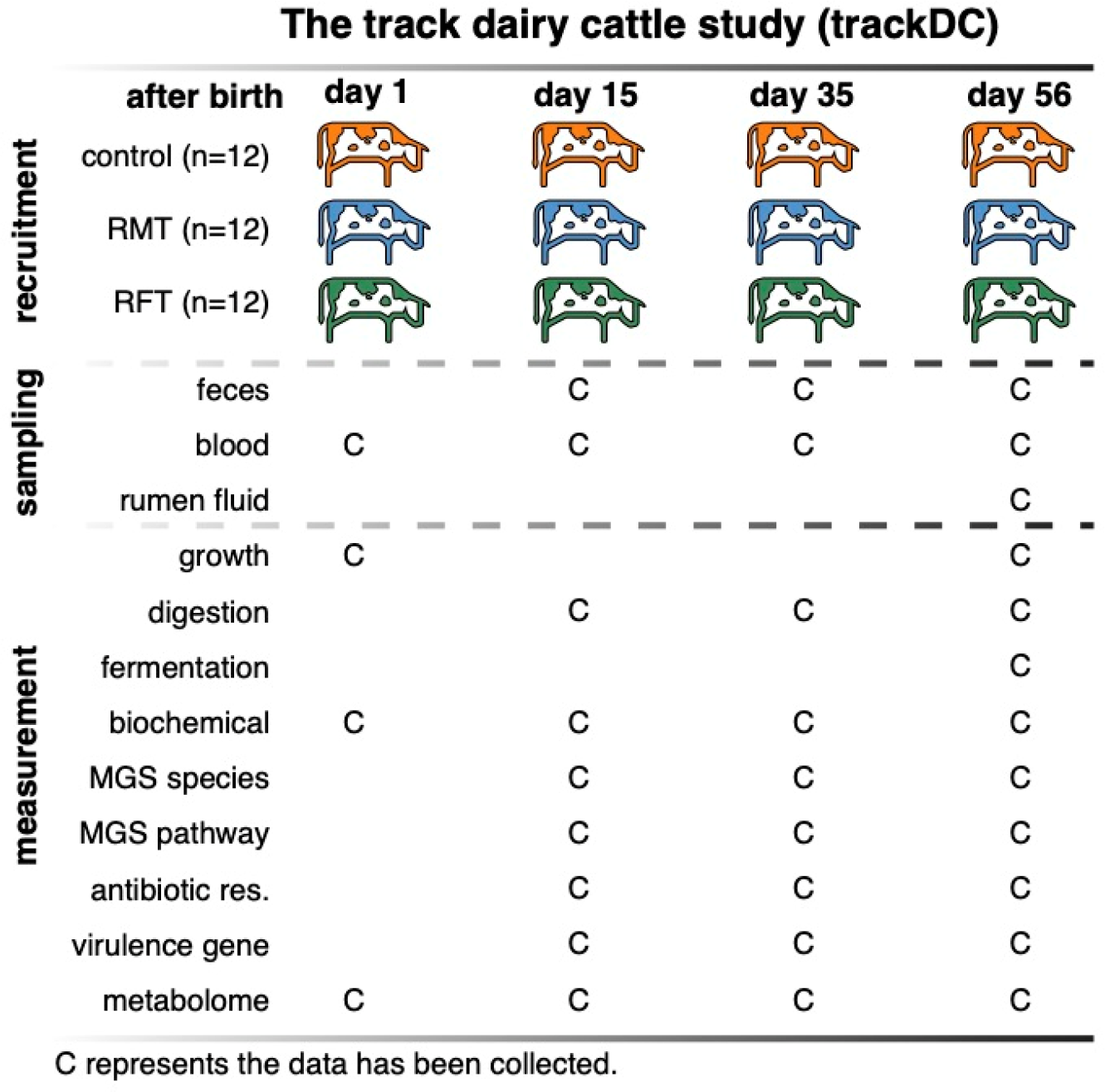
Study design and overview of the data that has been collected during the first two months.

**Figure S2.**
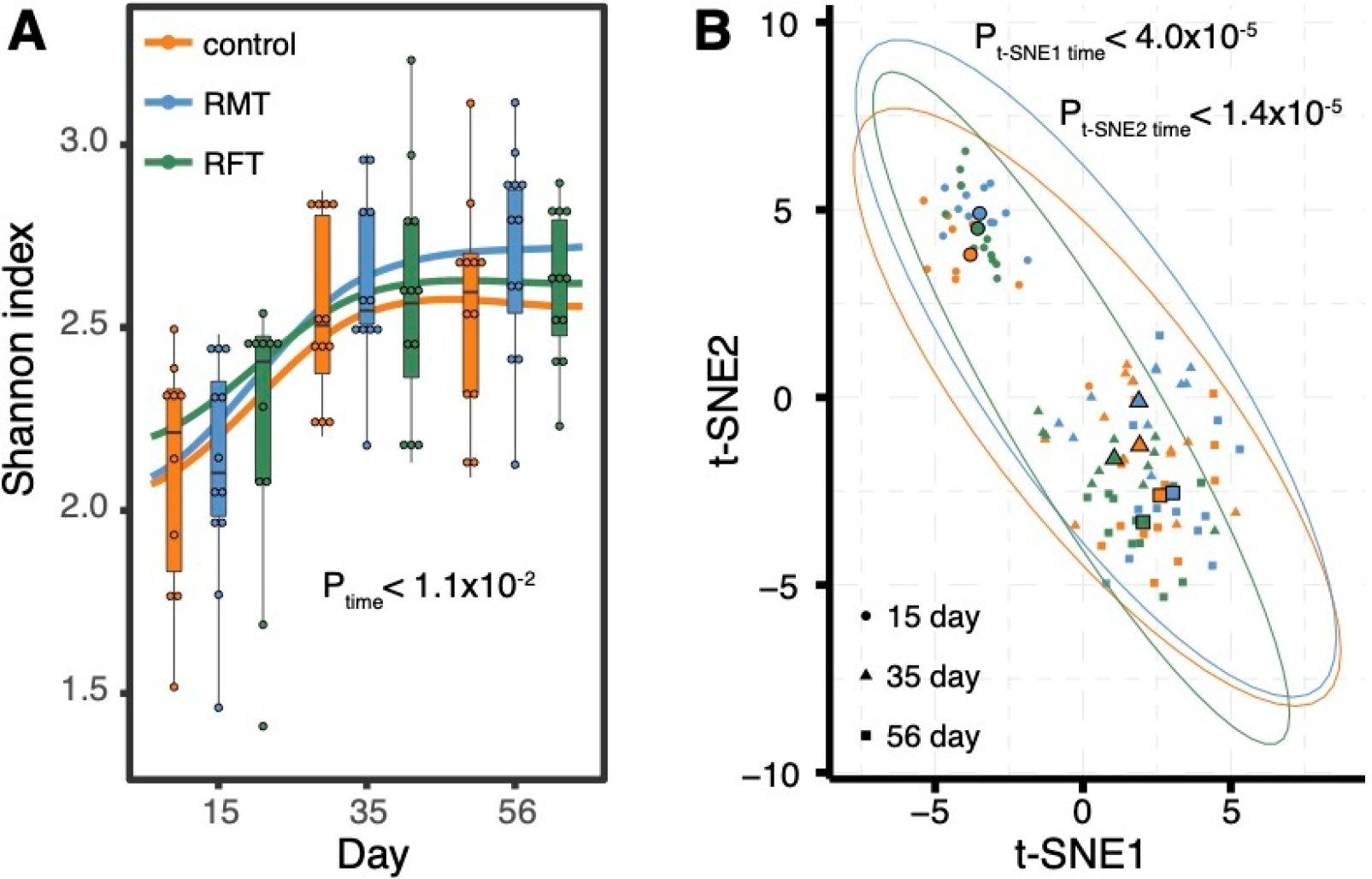
Temporal variations of the gut microbial diversity and composition in newborn calves. A. Temporal changes of the Shannon index based on species level abundance. P values from Kruskal test are shown accordingly. **B.** Temporal changes of the gut microbial composition on species level abundance. P values from Kruskal test are shown accordingly.

**Figure S3.**
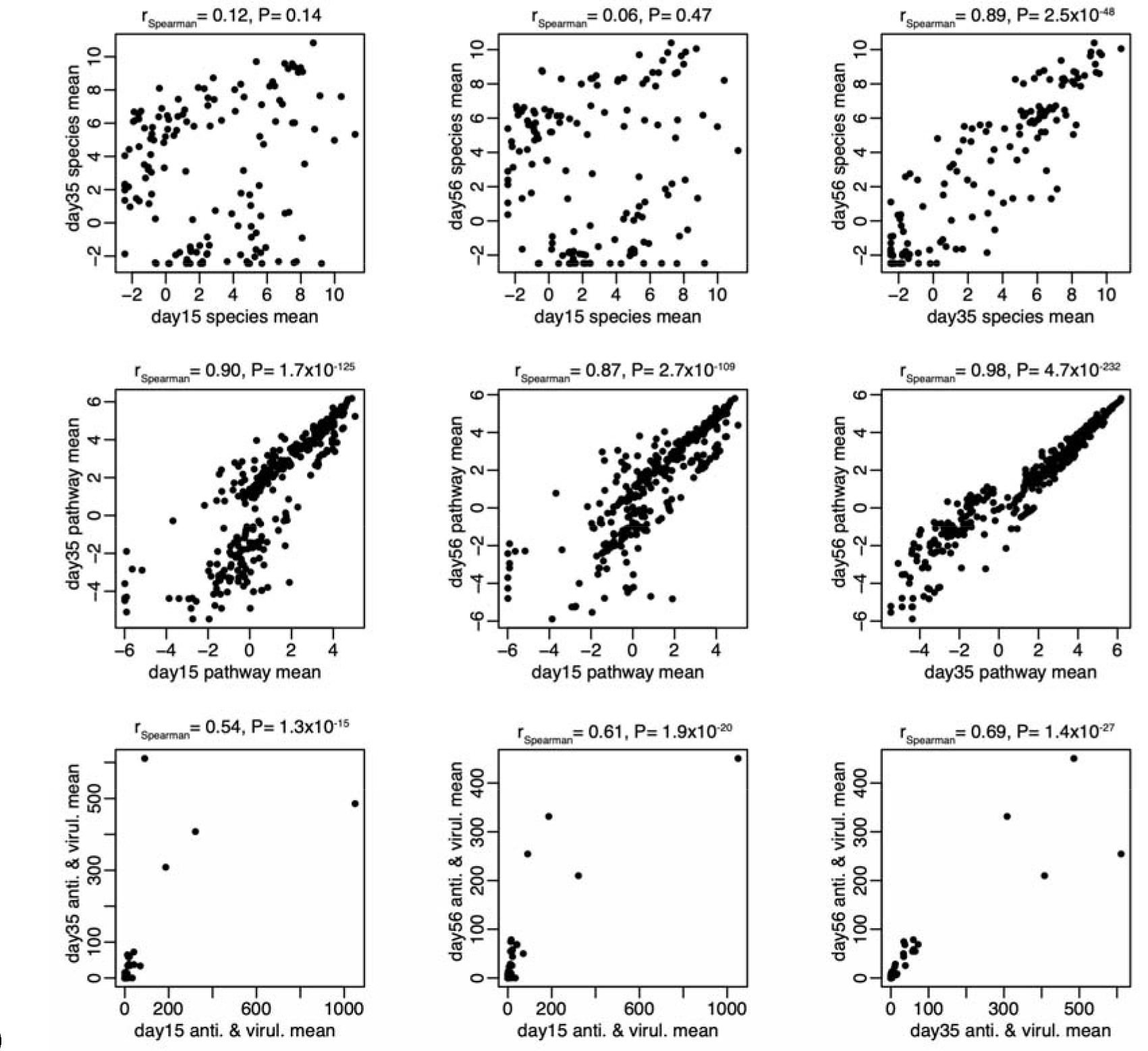
Comparison of microbial feature abundances between groups at different time points. Spearman correlation is applied to assess the associations of microbial abundances between different time points.

**Figure S4.**
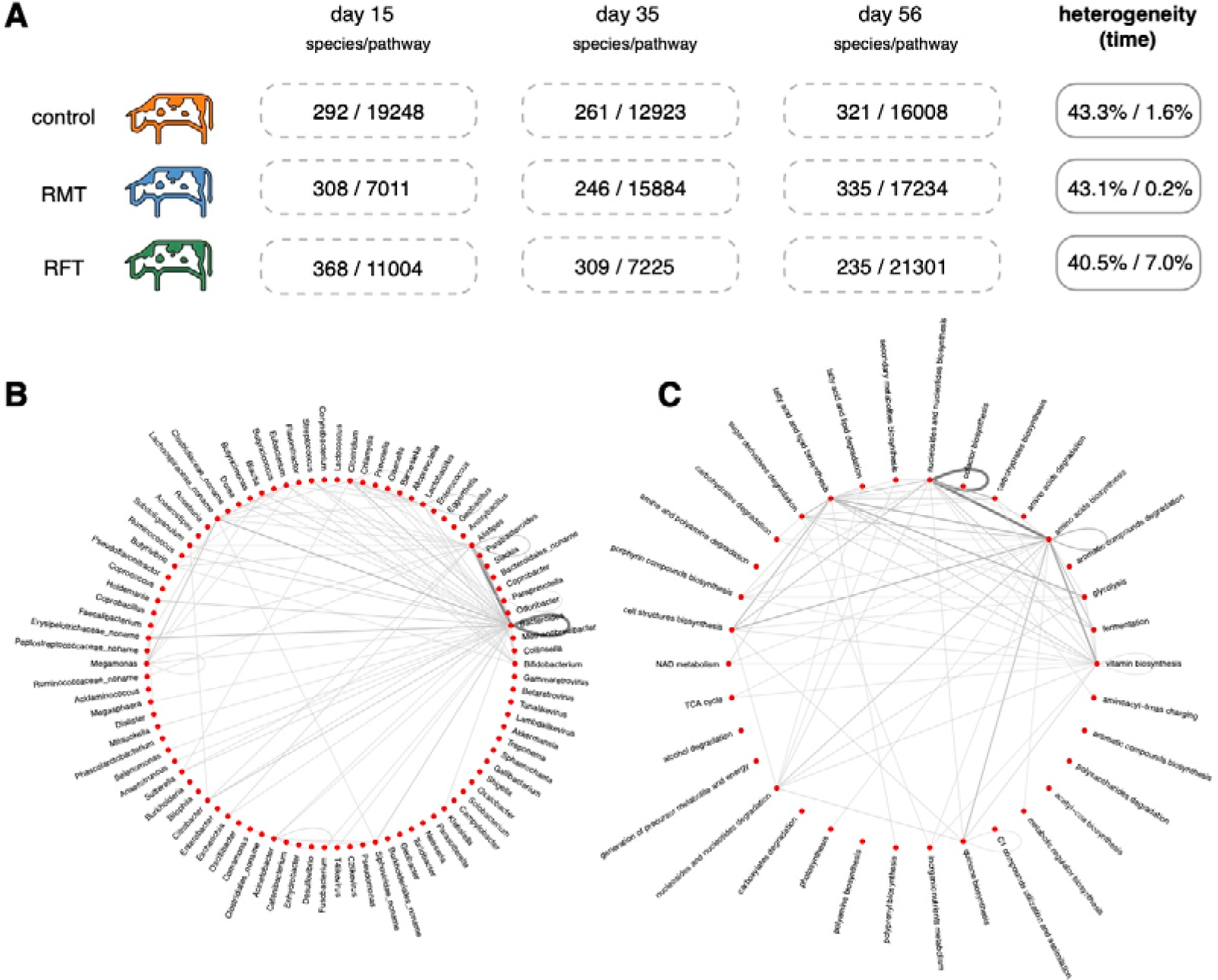
Temporal differences in microbial co-abundance networks. A. Summary of microbial species and pathway co-abundances in each group at different time points. The number of significant co-abundances has been listed and the heterogeneity of co-abundances is assessed by Cochran-Q test. **B.** Summary of differential microbial species co-abundances between time points. Each line represents differential species co-abundances between species from either the same or different genera. The width and darkness of the lines represent the relative number of differential co-abundances. **C.** Summary of differential microbial pathway co-abundances between time points. Each line represents differential pathway co-abundances between pathways from either the same or different metabolic categories. The width and darkness of the lines represent the relative number of differential co-abundances.

**Figure S5.**
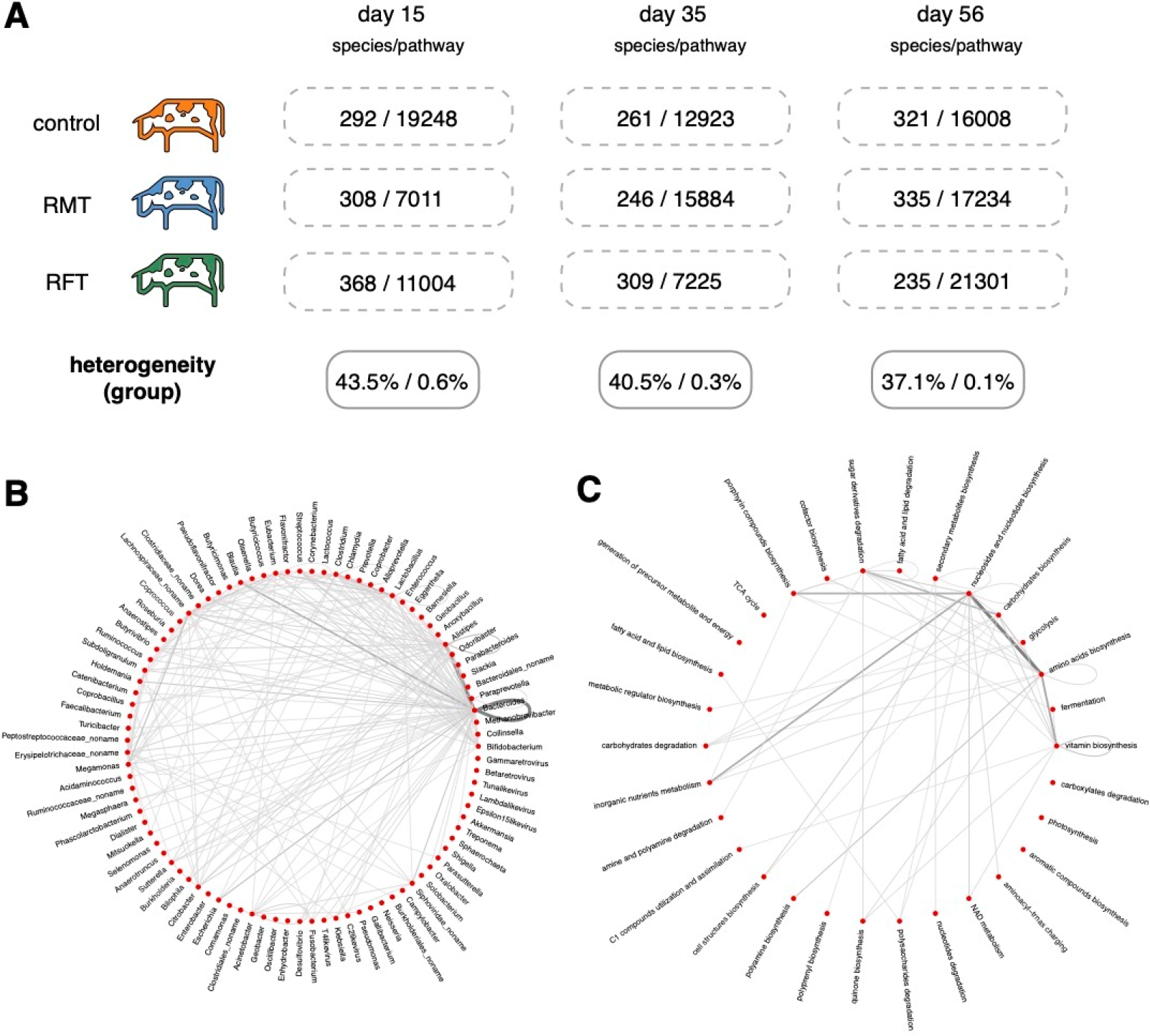
Differential microbial co-abundance networks between groups. A. Summary of microbial species and pathway co-abundances in each group at different time points. The number of significant co-abundances has been listed and the heterogeneity of co-abundances is assessed by Cochran-Q test. **B.** Summary of differential microbial species co-abundances between groups. Each line represents differential species co-abundances between species from either the same or different genera. The width and darkness of the lines represent the relative number of differential co-abundances. **C.** Summary of differential microbial pathway co-abundances between groups. Each line represents differential pathway co-abundances between pathways from either the same or different metabolic categories. The width and darkness of the lines represent the relative number of differential co-abundances.

**Figure S6.**
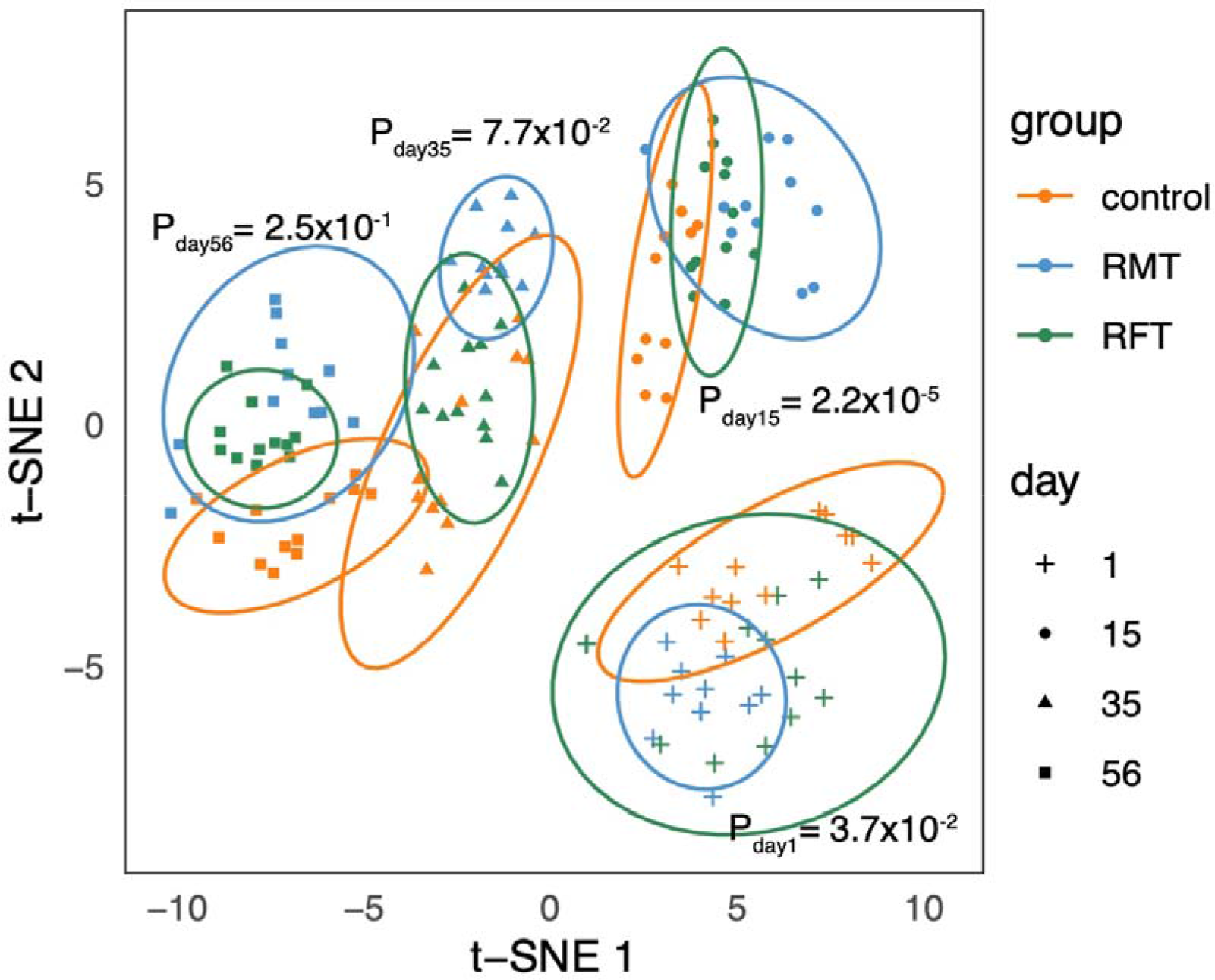
The composition of plasma metabolites shows temporal and group differences. P values from Kruskal test are shown accordingly

## TABLE LEGENDS

Table S1. Summary of phenotypes collected in this study

Table S2. Phenotypic differences between groups and time points

Table S3. Differential microbial species abundances between groups and time points

Table S4. Differential microbial pathway abundances between groups and time points

Table S5. Differential microbial antibiotic resistance and virulence gene abundances between groups and time points

Table S6. Microbial species co-abundances

Table S7. Microbial pathway co-abundances

Table S8. Microbial changes between day15 and day35 associated with phenotypic changes

Table S9. Microbial changes between day15 and day56 associated with phenotypic changes

Table S10. Summary of plasma metabolome

Table S11. Differential plasma metabolites between groups

Table S12. Microbial changes between day15 and day35 associated with metabolic changes

Table S13. Microbial changes between day15 and day56 associated with metabolic changes

Table S14. Metabolites mediate microbial impacts on phenotypic changes between day15 and day35

Table S15. Metabolites mediate microbial impacts on phenotypic changes between day15 and day56

